# Visually-evoked activity and variable modulation of auditory responses in the macaque inferior colliculus

**DOI:** 10.1101/2024.09.11.612507

**Authors:** Meredith N. Schmehl, Jesse L. Herche, Jennifer M. Groh

**Affiliations:** Department of Neurobiology, Duke University, Durham, NC 27708; Center for Cognitive Neuroscience, Duke University, Durham, NC 27708; Duke Institute for Brain Sciences, Duke University, Durham, NC 27708; Department of Psychology & Neuroscience, Duke University, Durham, NC 27708; Department of Biomedical Engineering, Duke University, Durham, NC 27708

**Author notes:** Corresponding Author: Jennifer M. Groh. The authors declare no competing financial interests.

## Abstract

How multisensory cues affect processing in early sensory brain areas is not well understood. The inferior colliculus (IC) is an early auditory structure that is visually responsive (Porter et al. 2007; Bulkin and Groh 2012a, 2012b), but little is known about how visual signals affect the IC’s auditory representation. We explored how visual cues affect both spiking and local field potential (LFP) activity in the IC of two monkeys performing a task involving saccades to auditory, visual, or combined audiovisual stimuli. We confirm that LFPs are sensitive to the onset of fixation lights as well as the onset of visual targets presented during steady fixation. The LFP waveforms evoked by combined audiovisual stimuli differed from those evoked by sounds alone. In single-unit spiking activity, responses were weak when visual stimuli were presented alone, but visual stimuli could modulate the activity evoked by sounds in a stronger way. Such modulations could involve either increases or decreases in activity, and whether increases or decreases were observed was variable and not obviously correlated with the responses evoked by visual or auditory stimuli alone. These findings indicate that visual stimuli shape the IC’s auditory representation in flexible ways that differ from those observed previously in multisensory areas.

**New & Noteworthy:** We find that the inferior colliculus, a primarily auditory brain area, displays distinct population-level responses to visual stimuli. We also find that visual cues can influence the auditory responses of individual neurons. Together, the results provide insight into how relatively early sensory areas may play a role in combining multiple sensory modalities to refine the perception of complex environments.

## Introduction

Visual and auditory processing streams were once thought to remain initially separate in the brain (Ghazanfar and Schroeder 2006; Schmehl and Groh 2021). Studies of how visual and auditory stimuli *interact* focused chiefly on late stages of processing: association areas such as intraparietal cortex or motor output stages such as the superior colliculus (SC) (e.g., (Witten and Knudsen 2005; Chandrasekaran 2017; Schmehl and Groh 2021)). At such late processing stages, two key features characterize response patterns: a) both modalities can lead to strong responses, i.e., spiking activity increases in response to either visual or auditory stimuli; and b) combinations of visual and auditory stimuli lead to greater increases in spiking activity than do either of the two stimuli when presented alone. Indeed, in the SC, combined visual and auditory stimuli are thought to lead to nearly linear summation of the responses evoked by each stimulus alone (particularly if the stimuli are suprathreshold) (Meredith and Stein 1986; Wallace et al. 1998; Stanford et al. 2005; Alvarado et al. 2007).

However, interactions between these sensory systems are now known to begin earlier than was previously appreciated, including at pre-cortical stages (Bulkin and Groh 2006; Ghazanfar and Schroeder 2006; Schmehl and Groh 2021). What *kinds* of interactions between visual and auditory signals occur in “early” areas is unknown, and may be different from those observed at later stages. Early sensory areas are typically generalists, playing a role in many aspects of sensory perception, whereas later areas such as the SC are thought to be focused on specific roles such as generating orienting movements of the eyes to the location of a sensory stimulus regardless of its modality (Ghazanfar and Schroeder 2006; Gandhi and Katnani 2011; Barrett 2012). And only some of those perceptual functions are likely to be under the influence of other sensory modalities. It is also not clear that sensory signals should necessarily add linearly; rather, a variety of different types of modulation may occur. Indeed, evidence for a diversity of types of interaction has been observed at the level of core auditory cortex (Schroeder and Foxe 2002; Ghazanfar 2005; Bizley et al. 2007; Kayser et al. 2007, 2008; Morrill and Hasenstaub 2018; Atilgan et al. 2018).

To gain insight into the properties of multimodal interactions in an even “earlier” sensory brain region, we turned to a predominantly auditory area, the inferior colliculus (IC). The IC is located two stops earlier on the auditory pathway than auditory cortex, to which it sends input via the medial geniculate nucleus as an intermediary (Huffman and Henson 1990; Winer and Schreiner 2005). Nearly all ascending auditory signals pass through the IC (Aitkin and Phillips 1984; Oliver 1984, 1987; Glendenning et al. 1992; Saint Marie et al. 1997; Winer and Schreiner 2005), making it necessary for hearing (Masterton et al. 1968; Thompson and Masterton 1978). Nevertheless, the IC also receives input from a variety of visually responsive areas, specifically visual cortex (Cooper and Young 1976), the superior colliculus (Harting 1977), and even the retina itself (Itaya and Van Hoesen 1982; Yamauchi and Yamadori 1982; Zhang 1984; Paloff et al. 1985). Projections from visually responsive areas have been found throughout the different subdivisions of the IC (Gruters and Groh 2012).

Visual responses in the IC were first identified by (Mascetti and Strozzi 1988) in the cat, and subsequently in monkeys by (Porter et al. 2007). As expected of an auditory structure, visual signals are known to be weaker and rarer than the auditory signals and differ substantially from classically visual areas in having unusual receptive field structures (Porter et al. 2007). However, they are present in all subdivisions of the IC (Bulkin and Groh 2012a), indicating their widespread importance.

Although an early report noted that visual stimuli could influence auditory responsiveness in the rat IC (Syka and Radil-weiss 1973), the question of how visual signals influence auditory responses has not yet been systematically evaluated in the primate IC. Here, we evaluated both single-unit and LFP signals in the monkey IC during an audiovisual localization task. We observed a discordance between visually-evoked activity in the LFP, which was robust, and spiking activity, where it was much weaker. Nevertheless, visual stimuli could alter auditory-evoked responses in both measures of activity. Intriguingly, the addition of a visual stimulus to an auditory stimulus could either increase or decrease the response, and the direction of the effect was not predictable from the unimodal responses. Together, these results indicate that the cross-modal interactions in this early auditory brain area are considerably different from those reported in the adjacent SC, suggesting that the computations supporting multisensory interactions may vary widely in the brain.

## Materials and Methods

### Subjects

Subjects were two rhesus macaques (*Macaca mulatta*), one female age 13 years (Monkey J) and one male age 4 years (Monkey D) at the start of training. All procedures were conducted at Duke University and approved by the Duke University Institutional Animal Care and Use Committee.

### Surgical Preparation

Each monkey underwent two stereotactic surgeries using isoflurane anesthesia and aseptic techniques, and post-surgical analgesics were administered. The first surgery served to implant a device to limit head motion during experiments. The second surgery consisted of implanting a recording cylinder (Crist Instruments, 2 cm in diameter) over a craniotomy to make both the right and left inferior colliculi (IC) accessible at an angle of 26 degrees from vertical in the coronal plane (approaching from the right for Monkey J and from the left for Monkey D).

### Equipment

During experiments, monkeys were seated comfortably in custom primate chairs (Crist Instruments) with heads restrained. The chair was situated in a dark single-walled sound attenuation chamber (Industrial Acoustics Company) lined with sound attenuating foam (Sonex). Visual stimuli were presented via green light-emitting diodes (LEDs), and sounds were presented from loudspeakers (Cambridge SoundWorks) calibrated to equal volume across speakers (see “Stimuli”). Monkeys performed a saccade task involving looking at the lights and/or sounds as described in detail below (see “Task”). Eye position was monitored using an Oculomatic Pro eye tracking camera and software (500 Hz sampling rate). On correctly-performed trials, juice rewards were delivered using a custom mouth piece and a PVC tube (ClearFLEX, Finger Lakes Extrusion) controlled with a solenoid valve (Parker Hannifin Corporation).

Neural recordings were conducted with 125 mm tungsten microelectrodes (FHC). A 19 mm diameter grid of holes (Crist Instruments) spaced 1 mm apart was placed into the recording cylinder to allow electrode penetrations, which were controlled by a microdrive (NAN Instruments LTD). Neural activity was collected with a Multichannel Acquisition Processor (MAP system, Plexon Inc.), passed through appropriate amplifiers, and sent to SortClient software for online spike sorting. Single unit spiking activity was extracted from signals bandpass filtered between 150 Hz and 8 kHz. Local field potential (LFP) signals were filtered from 0.7 to 300 Hz and stored at a sampling rate of 20 kHz (Plexon, Inc.). Spike times, eye positions, and task parameters were also stored for offline analysis (Plexon, Inc.; Ryklin Software, Inc.).

### Stimuli

As noted above, stimuli consisted of sounds and lights originating from loudspeakers and LEDs located on a horizontal row at the monkey’s eye level. Speakers and lights were located at -12 and +12 horizontal degrees in the monkey’s visual field. Sounds were bandpass filtered noise at center frequencies of 420 or 2000 Hz (11 kHz sampling rate, 55 dB SPL, ±100 Hz bandwidth, 10 ms on ramp). As detailed further under “Task,” the experiment involved only two locations and only two sound frequencies because the combinations of these conditions as well as the pairing with visual stimuli yielded a large number of combinations to be assessed for each neuron. The two locations were chosen to have one location in each hemifield, providing spatial balance for the saccade task. The two frequencies were chosen on the basis of previous studies of the monkey IC as likely to yield responses to at least one of the two sound frequencies (Bulkin and Groh 2011). Sounds were frozen for a given neuron, meaning a single bandpass stimulus was generated for each frequency at the beginning of the session and the same pattern was used throughout the session. Sounds were generated using custom software (Ryklin Software, Inc.). Lights were green light-emitting diodes (LEDs) placed on the front face of each speaker.

### Task

Monkeys performed tasks involving saccade targets at either one or two locations. The current paper reports results from the single-location trials; the two-location trials allowed us to explore the contribution of multisensory information to the processing of two simultaneous sounds (Caruso et al. 2018; Mohl et al. 2020; Glynn et al. 2021; Jun et al. 2022; Schmehl et al. 2024; Chen et al. 2024; Groh et al. 2024) and are the subject of a separate publication (Schmehl et al. 2025).

To initiate a trial, the monkey fixated on a central light (0 horizontal degrees from the midline, at eye level). After 600-700 ms, a visual, auditory, or combined audiovisual stimulus pair was presented at either -12 or 12 degrees while the fixation light remained on. After a further 600-800 ms of fixation, the fixation light was extinguished and the monkey had 500 ms to make a saccade to the target (which itself remained on until completion of the saccade; Figure 1). The reinforcement window around the target was 16 degrees wide by 60 degrees tall, to account for low precision in sound localization in the vertical domain (Jay and Sparks 1990; Gnadt et al. 1991; Blauert 1996; Barton and Sparks 2001). Monkeys then maintained fixation on the target for 200-250 ms to receive a fluid reward (typically Welch’s 100% grape juice diluted 30% in water). The additional two-location trials not analyzed here were very similar, except that two target locations were used simultaneously and the monkeys were required to make sequential saccades to both target locations (Schmehl et al. 2025). All conditions were randomly interleaved for each neuron.

**Figure 1:**
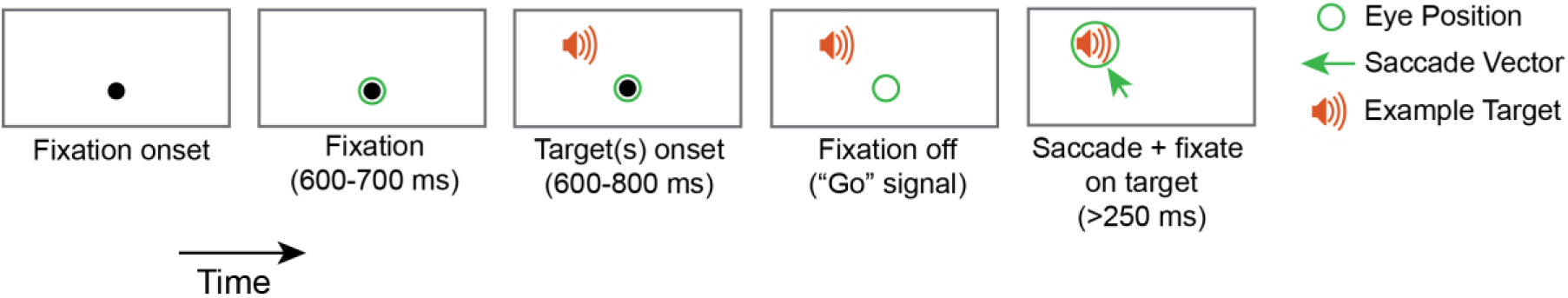
Task structure. Each trial began with the onset of a central light, and the monkey initiated the trial by fixating on the light for a variable period between 600 and 700 ms. A target was then presented (auditory, visual, or both at a single location), and the monkey waited 600-800 ms for fixation offset before making a saccade to the target (green arrow). After localizing the target, the monkey received a fluid reward. Trials continued in succession, and conditions were randomly interleaved.

Modalities were weighted such that 30% of the trials were visual, 30% of the trials were auditory, and 40% of the trials were audiovisual. Performance was >74% correct for both monkeys for all modalities, with performance on visual-only and audiovisual trials slightly exceeding that for auditory-only trials (Supplementary Figure 1). Table 1 shows a summary of the correctly-performed trials included for analysis.

**Table 1:** Number of trials in the dataset. Summary of correctly-performed trials that were included for analysis, grouped both by monkey and by modality. An additional 24,436 correct trials probed the responses to two simultaneous sounds; the results are reported separately (Schmehl et al. 2025).

|  | Visual | Auditory | Audiovisual | Total |
| --- | --- | --- | --- | --- |
| Monkey J | 5,141 | 4,429 | 6,981 | 16,551 |
| Monkey D | 2,994 | 2,522 | 3,813 | 9,329 |
| Total | 8,135 | 6,951 | 10,794 | 25,880 |

### Neural Recordings

The left and right IC in each monkey were localized using magnetic resonance imaging scans conducted at Duke University Hospital: sagittal localizing scan, coronal MPRAGE (T1 weighted 3D image, 0.5 mm), coronal T2 1.0 mm 55 slices with phase set right to left, coronal T2 0.5 mm 55 slices with phase set right to left (through the cylinder), and coronal T2 0.5 mm 55 slices with phase set anterior to posterior (through the cylinder). The location of the IC could be reconstructed accurately in reference to the coordinate system of a grid for holding electrodes in the recording cylinder (Crist Instruments). We recorded from the IC using a single tungsten microelectrode (FHC) while the monkey performed the task and neural data were collected as described above as well as previously (e.g., Willett & Groh, 2022). Neural data was recorded relative to a ground wire attached to the guide tube.

The IC of the monkey contains three physiologically-defined subregions: an anterior region in which neurons are relatively insensitive to sound frequency (the “untuned” region), a large region in which most neurons respond best to low frequencies, and a small central area in which frequency sensitivity progresses from low-preferring to high-preferring (Bulkin and Groh 2011). This tonotopic region is likely contained within the homologue to the central nucleus as defined in other species, but as it occupies only 17% of the total volume of the IC in monkeys (Bulkin and Groh 2011) and a nearly identical 18% in humans (Ress and Chandrasekaran 2013), a portion of the larger low-frequency preferring region of the monkey IC may also be consistent with the central-nucleus designation. For the purposes of the study concerning the two-location trials mentioned above, we were chiefly interested in sites where there was a difference between the responses to the low- and high-frequency stimuli (Schmehl et al. 2025). This ruled out neurons in the “untuned” region, but permitted inclusion of neurons from either the low-frequency or tonotopic regions. Of the 106 neurons recorded in this study, 13 (12.3%) responded best to the higher frequency sound and likely were from the tonotopic region. The remainder likely came from either the low-frequency region or the low-frequency tuned portions of the tonotopic region. Response latencies were consistent with known response latencies in the inferior colliculus (Ryan and Miller 1977, 1978; Groh et al. 2001; Maier and Groh 2010), with a median of 20 ms across the recorded population. The fastest latencies were about 15 ms. As is typical of long duration studies involving monkeys performing a behavioral task, reconstructing the locations of particular recording sites was not attempted.

### Data Analysis

All analyses were conducted on correctly performed trials, in which the monkey successfully localized the target (see Table 1). Only recording sites where the neuron was responsive to at least one modality (outlined below) and that had a minimum of 250 correct trials across both one-location and two-location conditions were included for analysis. This screen yielded an average of ∼244 correct one-location trials per neuron.

### Single Unit Activity

We recorded a total of 106 well-isolated single neurons, 82 from the right IC (Monkey J), and 24 from the left (Monkey D). Results were similar across monkeys and were combined for subsequent analysis.

Auditory responsiveness was assessed by comparing the activity during a time window after sound onset to a corresponding period of steady fixation immediately prior to sound onset. Two windows were used, either 0-100 ms or 0-500 ms after sound onset, to account for the presence of primarily transient vs. transient+sustained response patterns, which are known to occur in IC neurons (Ryan and Miller 1978; Willott and Urban 1978; Aitkin et al. 1994; Syka et al. 2000; Bulkin and Groh 2011). Units that exhibited a significant change in activity relative to baseline in either of these periods, or that passed visual inspection of peri-stimulus time histograms pooled across all auditory conditions, were considered auditory-responsive (pooled across all sound-only trials regardless of sound location or frequency, paired two-tailed t-test comparing firing rates before and after stimulus onset, p<0.025 for each tail, equivalent to p<0.05 for a one-tailed test). 106 neurons met these criteria.

Visual responsiveness was also assessed in several ways. Since each trial involved a visual fixation stimulus, activity was analyzed in a time window 50-250 ms after the onset of that fixation stimulus in comparison to a comparable baseline prior to fixation stimulus onset. This time window was chosen based on prior work, which showed that visual responses in the IC occur with a slight delay relative to stimulus onset (Gutfreund et al. 2002; Porter et al. 2007). The position of this fixation stimulus on the retina naturally varied based on where the eyes happened to be when the fixation stimulus appeared. Thus, we were able to conduct a rough evaluation of the receptive field for each neuron based on the location of the fixation light on the retina during passive looking at the time of fixation light onset. This was only an approximate assessment as it depended on uncontrolled variation in eye position prior to onset of the fixation light, which was low in this dataset because only a single fixation position was used.

We also examined responsiveness in a similar time period following entry of the eye into the fixation window, again in comparison to baseline (50-250 ms prior to fixation). Finally, on trials in which a visual saccade target was presented, we examined responsiveness 50-250 ms following target onset. As with the tests of auditory responsiveness, we used a paired two-tailed t-test, p<0.025 for each tail, deployed on all the trials involving visual stimuli regardless of their locations. Detailed results will be presented further below.

Visual influences on auditory responses were assessed differently. Activity was analyzed in a time window 0-500 ms after stimulus onset, to allow for visual influences that may occur throughout the trial. Because each sound could be presented either alone or with an accompanying visual stimulus, activity during each of the four auditory conditions (420 or 2000 Hz located at -12 or +12 degrees) was subtracted from the activity during the corresponding audiovisual condition. This difference was assessed separately for all audiovisual conditions in the dataset (4 conditions for each of 106 neurons, or 424 conditions in total). As with the tests of visual and auditory responsiveness, we used a paired two-tailed t-test, p<0.025 for each tail, to identify differences that diverged from zero, indicating a significant visual influence on auditory responsiveness.

### Local Field Potentials

Local field potential (LFP) signals were successfully analyzed from 83 of the 106 recording sites (the remainder were excluded due to technical problems affecting alignment of these data with the trial event data). As noted above, LFP signals were filtered from 0.7 to 300 Hz and stored at a sampling rate of 20 kHz (Plexon, Inc.), and were recorded with respect to a ground wire attached to the guide tube. Signals were normalized by subtracting the mean voltage during a 200 ms baseline period prior to fixation light onset across all trials for each recording site. Recording sites were included based on the significance of the associated single unit activity rather than LFP responsiveness.

LFP responsiveness was assessed in several ways, depending on the characteristics of the mean response across sites. For LFP signals aligned to fixation light onset (Figure 2a), we examined a 20 ms time window for each recording site and for each trial, centered at the positive peak of the mean response across sites (104 ms), in comparison to an equivalent time window prior to fixation light onset (centered at -104 ms). A recording site was deemed responsive if the distribution of the difference in the trial-wise mean LFP signals during these two periods was greater than zero (one-tailed t-test, p<0.05). For LFP signals aligned to visual target onset (Figure 2c), we examined two 20 ms time windows for each recording site and for each trial, centered at the positive peak (130 ms) and the negative peak (168 ms) of the mean response across sites, in comparison to an equivalent time window prior to visual target onset (centered at -130 and -168 ms). A recording site was deemed responsive if either time window passed the same test described above (one-tailed t-test, p<0.025 for each test, testing for a difference greater than zero for the positive peak and less than zero for the negative peak).

**Figure 2:**
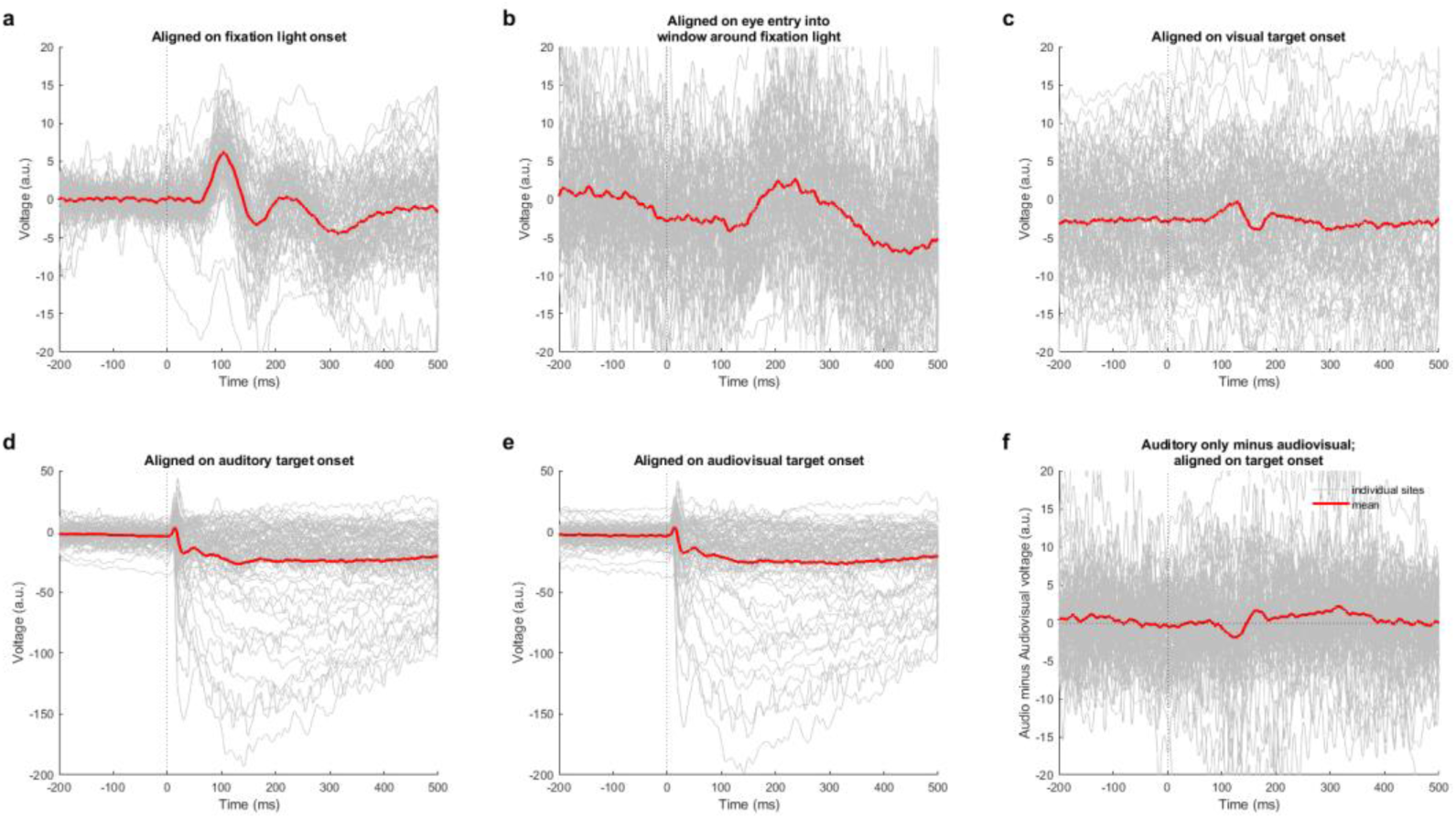
L**o**cal **field potentials display responses to fixation and target lights in addition to sounds.** Local field potential (LFP) aligned to various stages of the trial. a) LFP aligned on the onset of a fixation light. b) LFP aligned on the time of the monkey’s eye entry into a spatial window surrounding the fixation light. Note that sometimes the monkey anticipated the onset of the fixation light and no saccade was required; such trials are excluded from this analysis. c) LFP aligned on the onset of a visual target during steady fixation elsewhere. d) LFP aligned on the onset of an auditory target. e) LFP aligned on the onset of an audiovisual target. f) Difference in LFP shown in (d-e). In all subpanels, gray traces represent individual recording sites, and thick red traces represent the mean across all recording sites. Traces are averaged across all relevant stimulus conditions (e.g., traces aligned on auditory target onset in panel d include all sound frequencies and locations; those in e include all sound frequencies and locations that were paired with visual stimuli). Note the change in y-axis scale in (d-e) compared to (a-c), reflecting stronger auditory and audiovisual than visual responses. See Supplementary Figure 2 for a depiction of the mean traces accompanied by standard error for easier comparison to baseline responses.

Finally, for the difference in auditory and audiovisual target-aligned signals (Figure 2f), we examined two time periods informed by the shape of the mean response across sites. First, we examined a 20 ms time window for each recording site and for each trial, centered at the negative peak of the mean response across sites (122 ms), and tested whether the distributions of mean auditory and audiovisual trial-wise signals were different (one-tailed t-test, testing for a difference less than zero). Second, also for each recording site and for each trial, we examined a time window from the positive peak of the mean response (164 ms) to the end of the trial (500 ms), testing the difference in the distributions of mean auditory and audiovisual signals during this entire period (one-tailed t-test, testing for a difference greater than zero). A recording site was deemed responsive if either test indicated a significant difference from zero (p<0.025 for each test).

## Results

### Local Field Potential

Given that the IC is fundamentally an auditory structure, visually-evoked activity is expected to be weaker and rarer than auditory-evoked activity. Accordingly, the LFP is a particularly useful measure because it represents aggregate activity from a population of neurons, and can include pre- and post-synaptic activity in that population regardless of whether the threshold for triggering a spike was crossed. Therefore, we first probed the LFP, and we considered three points in time during the trial: the onset of the visual fixation light, the time of entry of the eyes into the reinforcement window around the fixation light, and the onset of the visual target (for trials with visual saccade targets).

Figure 2 shows LFP signals aligned to each of these time points. Figure 2a shows results aligned on fixation light onset for each recording site (individual gray traces). A distinct wave-like response occurs after fixation light onset for nearly every recording site. This wave exhibits a positive, then negative, peak before settling at a slightly negative plateau. We tested whether this positive peak differed from a corresponding period during the pre-fixation baseline and found that it did for 60 of 83 recording sites (72.3%) (one-tailed t-test, p<0.05; see Materials and Methods for details). In Figure 2b, the traces are aligned on the eye’s entry into the window around the fixation light, and a fresh visual response can be observed as the saccade brings the fixation light to a new (more foveal) retinal location. This response is most evident in the mean across sites (red trace). Finally, Figure 2c shows the responses to a visual target presented at either -12° or 12° during steady fixation (data are combined across both locations). These responses appear slightly weaker than those seen in connection to the onset of the visual fixation light. This could be because (a) they are added on top of any activity evoked by that target, so the deviation from the voltage level prior to target onset is modest, (b) they are presented from only 2 retinal locations, only 1 of which is in the contralateral hemifield, or (c) visually-evoked activity in the IC is context dependent and depends on the role or timing of the stimulus in the task. We will discuss these possibilities further in the Discussion. Overall, 19 of 83 recording sites (22.9%) exhibited either a significant positive or a significant negative peak in comparison to an equivalent baseline period (one-tailed t-test, p<0.025 for each test; see Materials and Methods for details).

In contrast, the sound-evoked LFP is quite strong. Figure 2d shows the signals on auditory trials (pooled across both frequencies and locations). Note the y-axis is adjusted to accommodate these much larger signals. Other than the greater magnitude, their character was similar to the visually-evoked LFPs, again usually beginning with a short-latency and short-duration positive peak, followed by a steadier and longer-lasting negative plateau. In other words, the peak sequences were similar, but occurred much faster on auditory than on visual trials.

The LFP evoked on audiovisual trials (again pooled across all stimulus combinations) is grossly similar to that observed on auditory-only trials (Figure 2e). However, when the mean audiovisual LFP for each site is subtracted from the corresponding auditory one (Figure 2f), a modest influence of the inclusion of the visual stimulus appears, particularly when averaged across sites (red trace deflects away from zero, first negative, then positive peaks). 54 of 83 recording sites (65.1%) exhibited a difference in the mean auditory and audiovisual LFP signals, defined as either a difference in the LFP signal at the time point of the mean negative peak, or a difference in the LFP signal between the time point of the mean positive peak and the end of the trial (one-tailed t-test, p<0.025 for each test; see Materials and Methods for details). This result indicates that 65.1% of these recording sites displayed a visual influence on auditory responsiveness.

Together, these findings reveal that visual signals are indeed reaching the IC in the context of this task, and that, as previously shown (Porter et al. 2007; Bulkin and Groh 2012a), the most obvious visual influences can be seen at the onset of the fixation light. Even so, the presentation of a visual target later in the trial does cause a measurable signal and a measurable modulation of auditory signals.

### Spiking Responses

We next considered individual neurons’ spiking responses to the onset of the fixation light, as previously reported by (Porter et al. 2007). Generally, these responses were quite weak. Figure 3a shows the individual (gray) and average peri-stimulus time histograms (PSTHs) (red) across sites. A few sites show distinct excitatory or inhibitory responses, but, in general, any visual effect on actual spiking was relatively uncommon in our dataset: the difference in the number of spikes observed 50-250 ms after fixation light onset was usually not very different from the number of spikes observed in the same window before fixation light onset (Figure 3b, Z-scores calculated by dividing the difference between the mean light-evoked and baseline firing rates by the standard deviation of baseline firing rates). In total, 8% of IC neurons showed significant excitatory responses (green bars) and 17% showed significant inhibitory responses (red bars; paired two-tailed t-test, p<0.025 for each tail). One possible explanation for the low prevalence of visual responses is that the visual receptive fields found in this population of IC neurons may be poorly probed by “chance” variation in eye position at the time of fixation stimulus onset. Figure 3c-f show the individual PSTHs (c,d) and receptive field maps (e,f) for the two most responsive neurons in the sample – while they are strongly responsive, their receptive fields are quite small and centered on the fovea. The fixation light only appeared at one location, and the monkeys’ eyes often lingered at or near that location while waiting for the next trial to start. This may have privileged detecting spike responses in a small set of foveally-sensitive IC neurons while disadvantaging the detection of spiking responses at other receptive field locations. The LFP signals may have been less sensitive to this limitation, by virtue of pooling over a larger population of neurons and/or inclusion of subthreshold signals.

**Figure 3:**
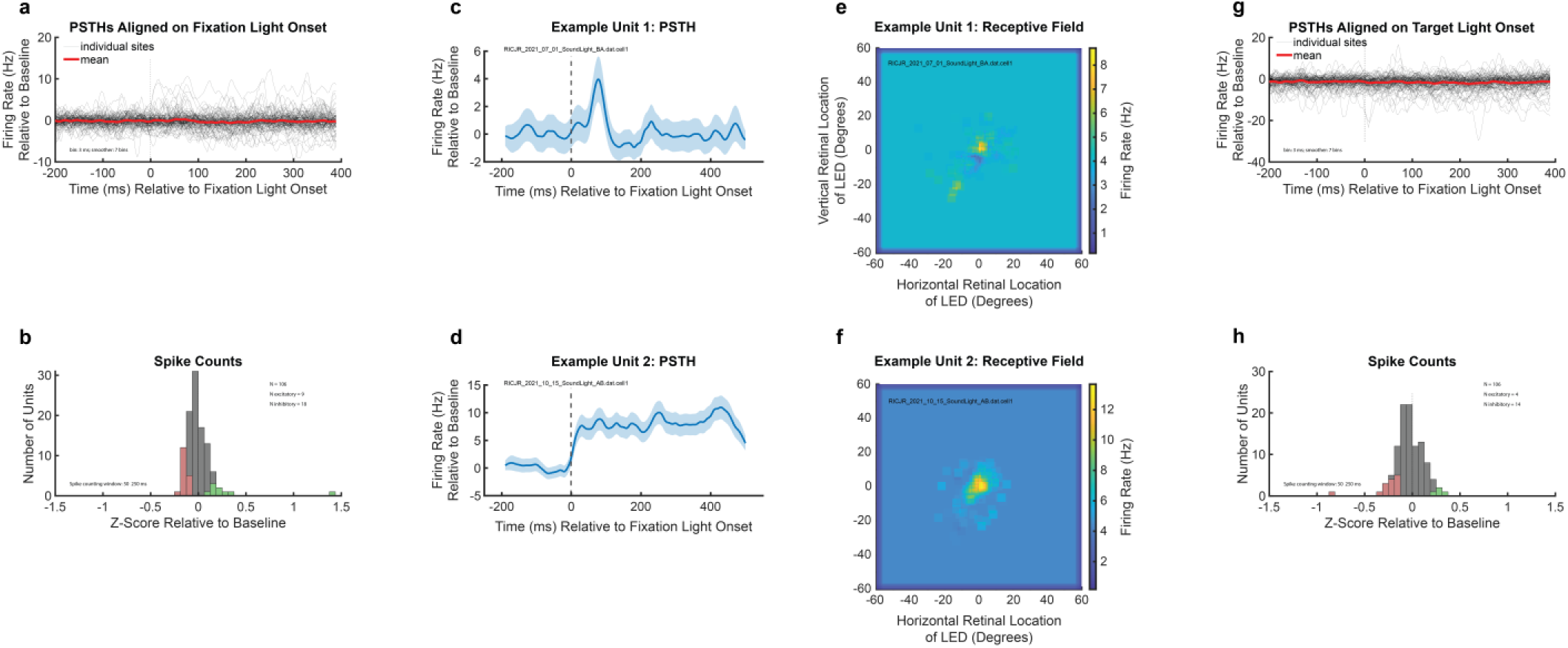
S**p**iking **activity in the IC reflects weak and rare responses to fixation and target lights during a localization task.** a) PSTHs of individual units (gray) and mean PSTH (red) for responses aligned to the onset of a fixation light presented at the horizontal midline and at eye level. b) Z-scores calculated as a difference between the mean light-evoked and baseline firing rates divided by the standard deviation of baseline firing rates. Green and red bars indicate units with a score significantly different from zero. c-d) Example responses of two individual neurons to the fixation light. Baseline firing rates were subtracted, and PSTHs were smoothed by convolving with a 5-point sliding triangle window and cubic spline interpolation. Shaded areas represent standard error. e-f) Spatial receptive fields of the same neurons as in (c-d), showing each neuron’s response to an LED fixation light falling on various horizontal and vertical locations on the retina during passive looking at the start of a trial. Baseline firing rates are indicated by the background color. Activity patterns were smoothed both horizontally and vertically using a 5-degree window. g) PSTHs of individual units (gray) and mean PSTH (red) for responses aligned to the onset of a target light. h) Z-scores calculated as a difference between the mean light-evoked and baseline firing rates divided by the standard deviation of baseline firing rates, using a spike counting window 50-250 ms after stimulus onset. Green and red bars indicate units with a score significantly different from zero. Results in panels g, and h are pooled across both visual target locations for individual neurons. Responses were also evaluated separately for the contralateral and ipsilateral targets (significant responses to the contralateral target: 4% excitatory, 4% inhibitory; for the ipsilateral target: 2% excitatory, 4% inhibitory; results not shown).

Similarly, Figure 3g shows the individual (gray) and average PSTHs (red) across sites for spiking responses to the onset of the target light during fixation. Like responses to the fixation light, these responses show visual effects on spiking activity in about 17% of the neurons (Figure 3h; 4% excitatory, green bars; 13% inhibitory, red bars; paired two-tailed t-test, p<0.025). In total, 37% of neurons exhibited significant responses in one or the other of these two time periods, with 6% responsive to both. When a neuron was responsive during both time periods, the direction of modulation (excitatory vs. inhibitory) was typically the same in these two cases. When separating out by target location, slightly more neurons were responsive to the contralateral stimulus (4% excited, 4% inhibited, or 8% overall) than to the ipsilateral target (2% excited, 4% inhibited, or 6% overall). (The lower proportion of responsive neurons when limiting to just one target location may be due to the impact of cutting the data in half on the odds of reaching statistical significance, i.e., reducing statistical power.) The prevalence of visual responsiveness was also similar in both monkeys: 23% of neurons in Monkey J and 33% of neurons in Monkey D were responsive to the fixation light, while 13% of neurons in Monkey J and 29% of neurons in Monkey D were responsive to the visual target. In summary, spiking modulation occurred in response to both fixation and target lights, but was rarer in this IC dataset in comparison to previous reports (Porter et al. 2007; Bulkin and Groh 2012a), perhaps due to the more sparse sampling of visual space.

We next asked how visual stimuli affect IC spiking activity when presented in combination with sounds. Specifically, we examined responses aligned to the onset of visual, auditory, and audiovisual targets following fixation onset during the task. Figure 4 shows the responses of four example neurons to visual (blue), auditory (orange), and audiovisual (purple) targets at single locations (pooled across frequencies and locations). Although these neurons were not responsive to visual input (blue), pairing that same visual cue with a sound (orange) produced a different auditory response (purple) – either by enhancing (Figure 4a) or suppressing (Figure 4b,c) the neuron’s auditory response. However, this was not always the case: Figure 4d shows an example of a neuron whose auditory responses appeared to be unaffected by the simultaneous presence of a visual stimulus.

**Figure 4:**
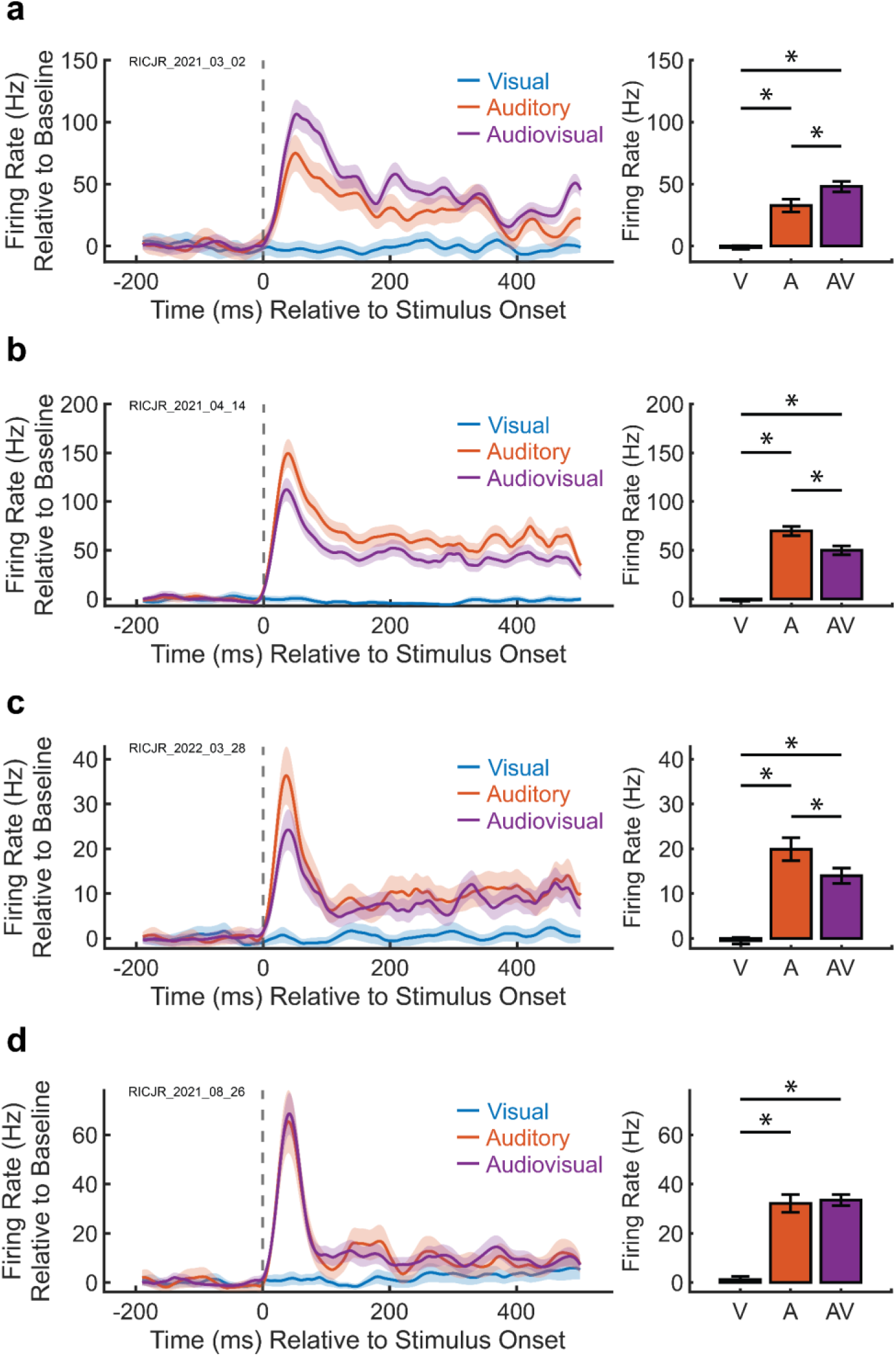
A visual cue can modulate a neuron’s auditory response. Example responses of four individual neurons to pooled combinations of all visual (blue), auditory (orange), and audiovisual (purple) conditions presented at a single location. Within each panel, the left subpanel is a peri-stimulus time histogram (PSTH) of each modality, and the right subpanel is a quantification of the average responses either in the first 500 ms after stimulus onset (0-500 ms) for sustained responses, or in the first 100 ms after stimulus onset (0-100 ms) for transient responses. Baseline firing rates were subtracted, and PSTHs were smoothed by convolving with a 5-point sliding triangle window and cubic spline interpolation. Shaded areas and error bars represent standard error. a) The presence of a visual cue augments the auditory response, and both the auditory and audiovisual responses are sustained throughout the stimulus presentation. b) The presence of a visual cue dampens the auditory response, and both the auditory and audiovisual responses are sustained throughout the stimulus presentation. c) The presence of a visual cue dampens the auditory response only immediately after stimulus onset, and the auditory and audiovisual responses are similar later in the stimulus presentation window. d) Auditory and audiovisual responses are similar, indicating no effect of the visual cue on auditory responsiveness. Note low visual responsiveness in all panels, indicating that the presence of a visual cue can modulate a neuron’s auditory response even if the neuron is unresponsive to visual input alone.

Given the low magnitude and low prevalence of visually-driven responses in these neurons (Figure 3), the visually-induced modulation of auditory responsiveness (Figure 4) is quite striking. These results suggest that although visual input may not drive a neuron’s spiking activity individually, pairing the same visual cue with a driving auditory input can be sufficient to modulate a neuron’s response, potentially to inform perceptual output.

We next quantified these effects at the population level. Figure 5a shows the differences between the responses to an audiovisual stimulus and the corresponding auditory stimulus alone, with each neuron contributing four data points for the four pairs of locations and frequencies in the task. About 15% of conditions showed a significant decrease in responsiveness (red/green bars, paired two-tailed t-test, p<0.025 for each tail), whereas 7.5% showed a significant increase in responsiveness.

**Figure 5:**
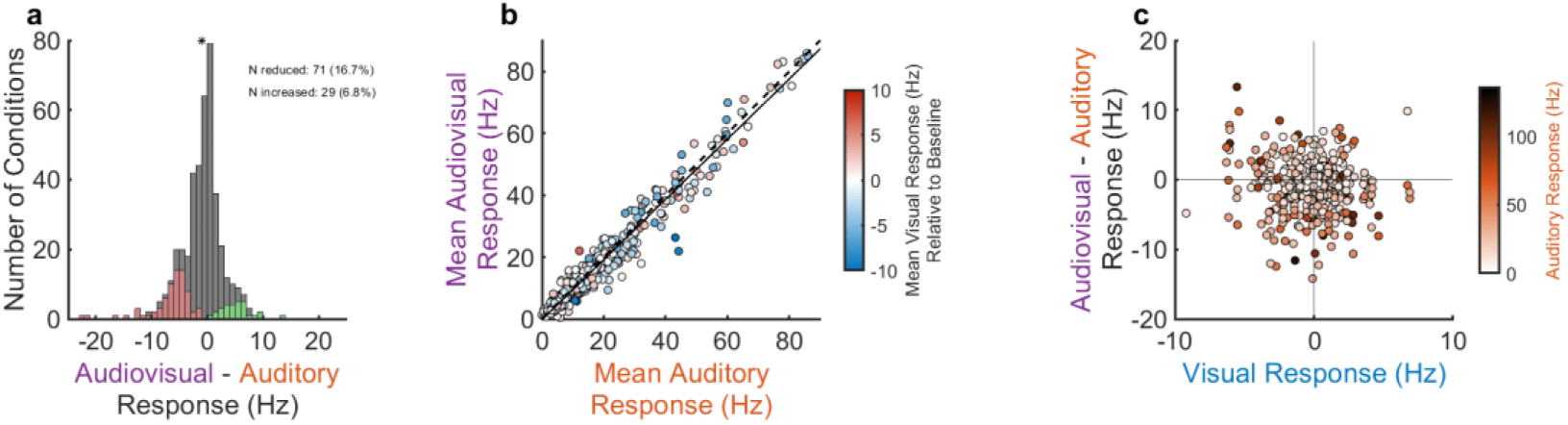
V**i**sual **influences on auditory responses are varied.** a) Distribution of audiovisual minus auditory responses, with each neuron contributing four conditions representing the four frequency-location pairs in the task. Visual stimuli added to sounds reduced activity in about 15% of conditions (red bars) whereas they increased the activity in about 7% of conditions (green bars; two-tailed t-test, p<0.025). In net, the population activity on audiovisual trials was slightly lower than on their auditory counterparts (p = 0.0383, paired sample t-test). Asterisk indicates distribution mean. Green and red bars indicate conditions with a value significantly different from zero. b) Comparison of mean auditory and audiovisual firing rates for each neuron, with colors indicating the corresponding visual firing rate for a visual cue presented at the same location. No strong relationship arises between visual firing rates and auditory or audiovisual firing rates, with both red and blue points falling both above and below the unity line. For clarity, visual responses are depicted relative to baseline, such that positive values indicate an excitatory response and negative values indicate an inhibitory one. No such baseline subtraction was applied to the auditory or audiovisual trials, but most were excitatory. c) Comparison of mean difference between audiovisual and auditory firing rates and baseline-subtracted visual firing rates, with colors indicating the corresponding auditory firing rate. Points arise in all four quadrants, indicating a wide degree of visual influences on auditory firing rates. In all subpanels, each neuron contributes 4 data points as sound frequencies and visual cue locations are counterbalanced (4 frequency-location pairs per neuron).

The direction of the effect was not strongly related to whether the visual response was excitatory on its own, nor was it strongly related to the magnitude of the auditory response on its own. Figure 5b shows the mean audiovisual response as a function of the mean auditory response, with color indicating whether the visual response was excitatory (red) or inhibitory (blue) relative to baseline. Both red and blue points lie both above and below the diagonal line of slope one, indicating that responses could be either increased or decreased when the visual and auditory stimuli were combined. Figure 5c shows the same pattern but from a different perspective – here the differences between the audiovisual and auditory responses are plotted as a function of the visual responses, with color indicating the magnitude of the auditory responses. Points lie in all 4 quadrants, indicating that both enhanced and reduced responses to audiovisual stimuli can occur regardless of whether the visual input is slightly excitatory or slightly inhibitory when presented on its own. A chi-squared test confirmed no significant relationship between visual excitation vs. inhibition and audiovisual enhancement vs. suppression (p = 0.196), indicating that all combinations of visual excitation/inhibition and audiovisual enhancement/suppression are common. Additionally, there is little evident relationship to the strength of the auditory response: darker red points lie in all four quadrants. These results further support the finding that the presence of a visual cue can modulate auditory responsiveness in the IC, but reveal that the direction of the effect – increasing vs. decreasing activity – can vary across the population and is not easily predicted from either the response to sound alone or the response to the visual stimulus alone.

## Discussion

Visual cues influence auditory perception at the behavioral level, but the role of early, predominantly unisensory areas in this process has remained relatively unexplored (for review, see (Ghazanfar and Schroeder 2006)). Here we investigated how visual cues modulate activity in the IC at population and single-unit scales and in response to visual stimuli presented alone or with sounds. We found that fixation lights and visual targets can evoke distinct wave-like responses in local field potential recordings, and that single units exhibit visual responses that are weaker but statistically evident in a modest subpopulation of neurons. Intriguingly, individual units’ responses to sounds can be modulated by the presence of a simultaneous visual cue, but in directions that are not easily accounted for by either the auditory or the visual responses alone. Indeed, modulation of auditory signals could be seen even when the visual stimulus did not itself evoke an overt spiking response. This suggests a complex form of interaction between visual and auditory signals in this nominally auditory structure.

These findings extend previous work that identified visual and eye movement-related responses (Porter et al. 2007; Bulkin and Groh 2012a) and eye position-dependent auditory responses (Groh et al. 2001; Zwiers et al. 2004; Porter et al. 2006) in the IC. The chief difference between our current study and the earlier studies from our group (Porter et al. 2007; Bulkin and Groh 2012a) is that the driven visual responses appeared to be less prevalent here. It is not yet clear what may account for this difference. One possibility is that the current study sampled visual space in a more limited way – there was only a single fixation location and only two visual stimulus locations, limiting the sampling of retinal receptive fields. Another possible factor is that the recordings in the current study were restricted to the low-frequency and tonotopic regions within the IC, whereas previous work intentionally sampled a wide range of locations spanning a larger portion of the IC (Bulkin and Groh 2012a). The difference in the variety of recording sites sampled may mean that the recording sites in the current study had a lower overall degree of visual input than in prior studies. It is known that the IC’s subregions differ in the prevalence of visual receptive fields with lower incidence in the low frequency-responsive and tonotopic regions that we tested here (Bulkin and Groh 2012a). Other attributes of the stimuli, tasks, or research subjects may also contribute to differences in visual responsiveness across studies.

Our results are similar to those in auditory cortex, where a variety of multisensory interactions have been identified. In auditory cortex, visual cues can elicit their own responses as well as modulate auditory responses in both excitatory and inhibitory ways (Bizley et al. 2007; Kayser et al. 2007, 2008; Morrill and Hasenstaub 2018). Non-auditory cues can also modulate current source density in auditory cortex, either by eliciting direct visual responses (Schroeder and Foxe 2002) or by modulating an auditory response by resetting the phase of neural oscillations (Lakatos et al. 2007). These effects are also evident for stimuli with high behavioral relevance; images or videos of faces can elicit their own independent responses as well as modulate LFP activity in response to vocalizations (Ghazanfar 2005). Finally, visual stimuli can modulate auditory responsiveness in more complicated ways, such as influencing which of two sounds is predominantly represented by individual neurons or the neural population (Atilgan et al. 2018).

Together, our results and those of these other prior studies in auditory cortex differ from related investigations in more multimodal areas such as the superior colliculus (SC). In the SC, which is reciprocally connected to the IC (Woollard and Harpman 1940; Moore and Goldberg 1963; Powell and Hatton 1969; Harting 1977; Adams 1980; Covey et al. 1987; Coleman and Clerici 1987; Sparks and Hartwich-Young 1989; Doubell et al. 2000; Hyde and Knudsen 2000; Caño et al. 2006; Stitt et al. 2015), spiking responses to both visual and auditory stimuli are common. Combinations of visual and auditory stimuli are generally thought to increase the activity of individual neurons, most often in an additive fashion (Meredith and Stein 1986; Wallace et al. 1998; Stanford et al. 2005; Alvarado et al. 2007). In contrast, audiovisual responses in the IC can be either slightly higher or slightly lower than auditory-alone responses. These patterns occurred even when a neuron had no visually-driven response, and rarely appeared to be additive.

A likely factor in these differences is that IC is thought of as a general-purpose auditory area, in contrast to the more overtly multimodal/oculomotor SC. The IC is located only a few synapses along the auditory pathway from the cochlea, and nearly all ascending auditory input passes through this brain region (Aitkin and Phillips 1984; Oliver 1984, 1987; Glendenning et al. 1992; Saint Marie et al. 1997; Winer and Schreiner 2005). Thus, it is likely that the overall purpose of this structure is to process sound, and that any visual signals in this area serve a purpose of modulating those responses rather than simply adding to them. Such modulation is likely to take multiple forms – perhaps even changing the spatial or frequency sensitivity of a neuron’s auditory signals – and not reduce easily to a simple monotonic (whether additive, subadditive, or superadditive) operation in which stronger input signals lead to more output (spiking) activity.

The specific mechanisms for such modulation may involve synaptic potentials that do not always lead to spiking activity, potentially explaining why visual signals are more evident in the LFP, which represents aggregate neural signals, than in the single unit responses, which derive only from spikes. Such synaptic potentials could lead the individual neurons to respond either more or less strongly to their auditory inputs, and could exert their effect not directly on the individual neuron being recorded from but on inputs to that neuron from adjacent neurons in the population. If the effects are indirect, this could explain differences between the observed overt visual responses and the net impact of combining visual and auditory stimuli in spiking activity – a neuron could show an excitatory response to a visual-only stimulus but a reduced response to sounds paired with visual stimuli if the latter case involves additional synaptic interactions and thus changes in responses upstream from the neuron being recorded from.

Our study provides a first look at the scope of the IC’s visual-auditory interactions and the factors that govern them. More work will be needed to explore such factors. For example, full mapping of visual receptive fields should be combined with assessing the role of visual-auditory spatial alignment on IC activity. This is a particularly interesting unexplored domain because the native codes for visual and auditory space are so different. The optics of the eye cause light from different locations in the environment to land at particular locations on the retinal surface, leading neurons throughout the visual pathway to have circumscribed receptive fields. In contrast, sound location is computed in part from binaural difference cues, which are proportional to the horizontal eccentricity of a sound. Neurons in the primate IC (and SC) do not exhibit circumscribed receptive fields, but rather appear to encode auditory space as a meter or rate code in which a large population of neurons responds with a particular level of activity, depending on where a sound is located along the azimuth (primate IC: Groh et al., 2003; Zwiers et al., 2004; primate SC: Lee & Groh, 2014; see also: Middlebrooks et al., 1994, 1998; McAlpine & Grothe, 2003; Werner-Reiss & Groh, 2008; Salminen et al., 2009). The brain must combine these disparate visual and auditory codes to understand where visual and auditory signals arise in space, and the IC is emerging as a site where such combination may occur.

In addition, further work is needed in the frequency domain. For example, at the behavioral level, visual lip reading cues can modulate the perception of speech sounds (e.g., the McGurk effect, (Sams et al. 1991; Wright et al. 2003; Ghazanfar 2005; Kayser et al. 2007, 2008; Hoffman et al. 2008; Nath and Beauchamp 2012)). This perceptual phenomenon might emerge from visually-induced changes in the representation of sound frequency within the auditory pathway. Further exploration is needed to ascertain the properties that govern these neural responses and their perceptual correlates.

Another key difference between our study and much of the work quantifying visual-auditory interactions in the superior colliculus is the task and behavioral state of the animal. Here, monkeys were awake and performing a localization task involving saccades to the respective stimuli. Much of the early work detailing the properties of multisensory interactions in the superior colliculus was conducted in anesthetized animals (Meredith and Stein 1986; Stanford et al. 2005). Anesthesia is known to affect the signals in the SC (Populin 2005), and performance of a behavioral task is known to affect auditory signals in both the SC and IC (IC: (Ryan and Miller 1977; Metzger et al. 2006); SC: (Jay and Sparks 1984)). Behavioral state may have a particularly strong impact on signals that are near threshold for spiking, as was found here. Thus, a detailed comparison of multimodal interactions in the IC and SC will require use of common methods in both structures.

More work will be needed to refine our understanding of specialization of function within the IC and potentially in its interactions with other auditory structures. In many species, the IC is thought to contain a central nucleus and one or more surrounding regions (for review, see (Winer and Schreiner 2005)). These regions differ in their auditory connection patterns, but all receive input from multiple modalities – specifically visual, somatosensory, and oculomotor signals (for review, see (Gruters and Groh 2012)). However, while anatomical inputs exist that can in principle mediate visual responses, the strength of responsiveness to visual signals doesn’t appear to be uniform: in a previous mapping study, visual signals appeared weakest in the deepest sites of the IC that are most likely to lie within the central nucleus (Bulkin and Groh 2012a). Thus, there may be differences in how non-auditory and auditory signals interact in the different subregions of the IC. A study geared toward reconstructing the recording sites in relation to these subregions will be needed to answer this question.

Collectively, these experiments provide insight into the influence of visual cues on auditory perception. Together with the other studies in related auditory regions, the findings presented here shed new light on the connections between multiple sensory systems and the ways in which context from another modality can inform processing of a brain region’s “dominant” modality. As the field of neuroscience moves forward, it will be critical to consider how sensory systems interact to give rise to complex behaviors in varied sensory environments.

## Supplementary Materials

**Supplementary Figure 1:**
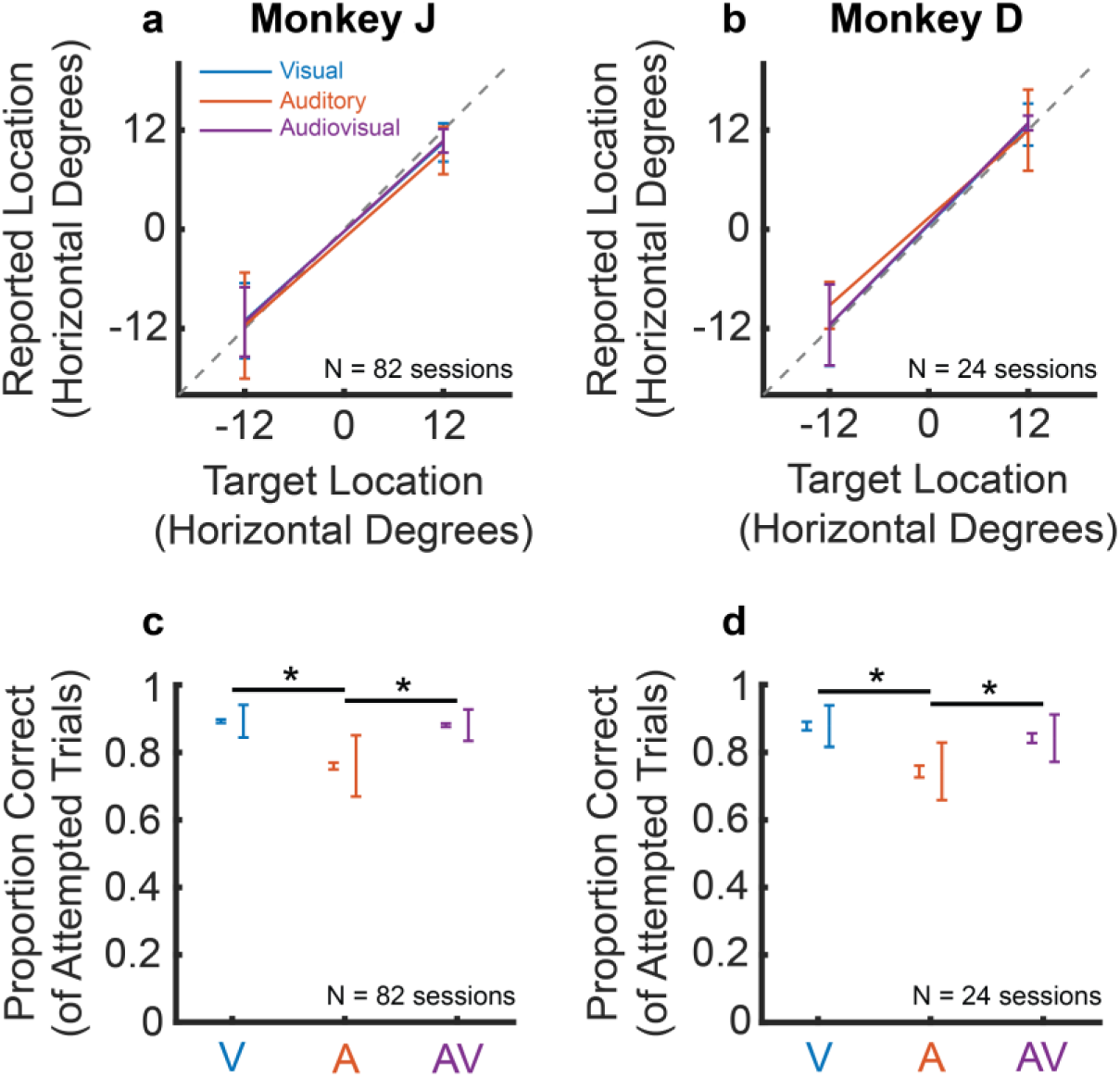
Monkeys accurately perform an audiovisual localization task. a-b) Mean and standard deviation of saccade endpoints during trials performed by each monkey for visual, auditory, and audiovisual stimuli presented at the two horizontal target locations in the task. Endpoints for all three modalities are largely overlapping and are aligned with the actual target location. c-d) Proportion of attempted trials of each modality in which each monkey correctly localized the targets within a 16-degree wide by 60-degree tall window. Visual and audiovisual conditions had a higher proportion of correct trials than auditory conditions, but all types of conditions were performed at an average accuracy of approximately 74% or greater. Two sets of error bars are shown: left error bars represent standard error and right error bars represent standard deviation, calculated across the number of sessions for each monkey.

**Supplementary Figure 2:**
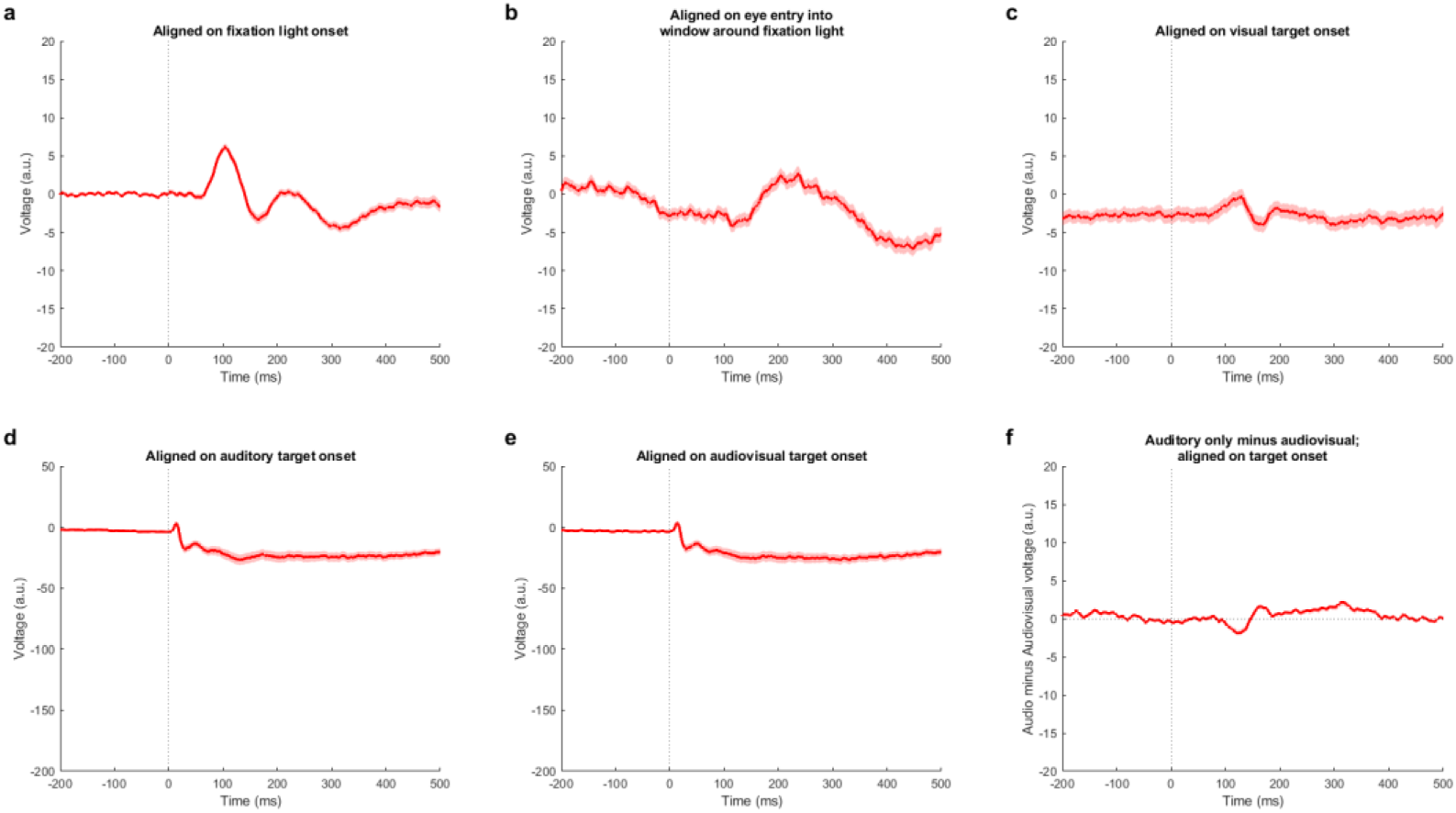
Mean local field potentials diverge from baseline in responses to fixation lights, target lights, and sounds. Same mean local field potential (LFP) responses as shown in Figure 2, with shaded areas representing standard error. As in Figure 2, responses are aligned to various stages of the trial. Standard error remains narrow and diverges from baseline in all cases.

